# Dynamics of myogenic differentiation using a novel Myogenin knock-in reporter mouse

**DOI:** 10.1101/2020.12.21.423736

**Authors:** Maria Benavente-Diaz, Glenda Comai, Daniela Di Girolamo, Francina Langa, Shahragim Tajbakhsh

## Abstract

**Background:** *Myogenin* is a transcription factor that is expressed during terminal myoblast differentiation in embryonic development and adult muscle regeneration. Investigation of this cell state transition has been hampered by the lack of a sensitive reporter to dynamically track cells during differentiation.

**Results:** Here, we report a knock-in mouse line expressing the tdTOMATO fluorescent protein from the endogenous *Myogenin* locus. Expression of tdTOMATO in *Myog*^*ntdTom*^ mice recapitulated endogenous *Myogenin* expression during embryonic muscle formation and adult regeneration and enabled the isolation of the Myogenin^*+*^ cell population. We also show that tdTOMATO fluorescence allows tracking of differentiating myoblasts *in vitro* and by intravital imaging *in vivo*. Lastly, we monitored by live imaging the cell division dynamics of differentiating myoblasts in vitro and showed that a fraction of the MYOGENIN^+^ population can undergo one round of cell division, albeit at a much lower frequency than MYOGENIN^-^ myoblasts.

**Conclusions:** We expect that this reporter mouse will be a valuable resource for researchers investigating skeletal muscle biology in developmental and adult contexts.

## BACKGROUND

Embryonic and postnatal myogenesis as well as adult muscle regeneration are regulated by a family of basic helix-loop-helix myogenic regulatory factors (MRFs) comprising *Myf5, Mrf4, Myod* and *Myogenin* (*Myog*). Following myogenic specification in the embryo, the MRFs are expressed in a sequential manner to ensure commitment, proliferation, differentiation, and fusion to give rise to multinucleated skeletal myofibres. Single and combinatorial mouse knockout models of the MRFs have established a genetic hierarchy where *Myf5, Mrf4* and *Myod* control lineage commitment and proliferation of myogenic progenitors and *Myod, Mrf4* and *Myog* are involved in terminal differentiation [1]. Notably, among the single MRF knockout mice, only *Myog-*null homozygous animals die at birth due to severe skeletal muscle defects [1–3]. Thus, unlike the other myogenic bHLH factors, *Myog* has no redundant or compensatory mechanisms to replace its function during development. Myoblasts lacking this gene accumulate in the muscle-forming areas throughout the body and fail to form normal myofibers in vivo, pointing to its critical role in terminal differentiation of myoblasts (Hasty et al., 1993; Nabeshima et al., 1993; Venuti et al., 1995). While in *Myog*-null embryos have some disorganized residual primary fibres, major differences between mutant and wild-type embryos become apparent during the initiation of secondary myofibre formation [2,4]. Unexpectedly, conditional ablation of *Myog* during the perinatal and postnatal period does not result in noticeable defects in muscle morphology or histology, suggesting that *Myog*^-/-^ myoblasts can still contribute to muscle growth [5,6]. Additionally, conditional ablation of *Myog* in a Duchenne muscular dystrophy mouse model (*mdx* [7]) did not result in an adverse phenotype, confirming that *Myog* is dispensable for adult muscle regeneration [8]. Nevertheless, although *Myog*-null muscle stem cells (MuSCs) proliferate and differentiate in culture as efficiently as wild-type cells, the muscle gene expression program is profoundly altered in the absence of *Myog* (Meadows et al., 2008).

Adult muscle regeneration depends on MuSCs, characterised by the expression of *Pax7* [9–13]. Upon muscle injury, MuSCs activate the expression of *Myod*, proliferate to generate myoblasts that differentiate and fuse to form myofibres. Different reporter mouse lines have been generated to fluorescently label the *Pax7*^+^ muscle progenitor population, either from the endogenous locus [14–16] or as transgenes [15,17,18] thereby allowing imaging and isolation of *Pax7*-expressing cells. Additionally, inducible reporters in which expression of the *Cre* recombinase under the control of the *Pax7* promoter recombines a membrane or cytoplasmic fluorophore [13,19,20] have been used for permanent marking the myogenic lineage [21–23] and for live imaging [24]. Although several reporter mouse lines have been generated to identify differentiating myoblasts based on expression of *Myosin light chain* [25], *Myog* [26–28] and *Muscle creatine kinase* [29]), they are based on *lacZ* (*β-galactosidase* activity, [30]) or *cat* (*chloramphenicol acetyltransferase*, [31]) expression and thus only allow endpoint measurements on fixed samples.

Terminal myoblast differentiation is characterised by the expression of *Myog*, the cyclin-dependent kinase inhibitor *p21* and cell cycle withdrawal [32–34]. Experiments using the nucleotide analogue BrdU have shown that MYOG positive cells can undergo DNA replication [32], but it is still unclear how many divisions they can execute before definitively leaving the cell cycle.

Here, we took advantage of the CRISPR/Cas9 system, which allows precise genome editing [35],to generate a knock-in mouse line expressing a nuclear localised tandem-dimer Tomato (tdTOM) protein under the control of the endogenous *Myog* promoter, while retaining expression of MYOG protein. We show that heterozygous *Myog*^*ntdTom*^ mice exhibit robust reporter gene expression in fixed and live myogenic cells thus allowing *in vitro* and intravital microscopy studies of the dynamics of muscle differentiation and cell cycle withdrawal.

## MATERIALS AND METHODS

### Mouse maintenance

Animals were handled according to national and European Community guidelines and an ethics committee of the Institut Pasteur (CETEA, Comité d’Ethique en Expérimentation Animale) in France approved protocols (Licence 2015-0008). Except when indicated otherwise, males and females of 2-4 months were used.

### Generation of the Myog-ntdTomato construct for CRISPR-Cas9 mediated homologous recombination

A fragment of 1000 bp from the last exon of *Myog* was amplified by PCR from murine gDNA (primers 1 and 2, **Supplementary Table 3**), introducing SalI and NotI restriction sites. This fragment was subcloned into the donor plasmid encoding for tdTOM (kind gift from Dr. Festuccia). A fragment of 760 bp from the 3’UTR of the *Myog* gene just after the STOP codon was amplified by PCR from murine gDNA (primers 3 and 4). This amplification also introduced a mutation in the PAM sequence necessary for CRISPR-Cas9 genome editing. Using the PacI and SpeI restriction sites added, the fragment was subcloned into the PacI and XbaI digested tdTOM plasmid. Oligos containing a T2A (primers 5 and 6) [36] peptide and a triple NLS sequence from SV40 large T [37] were annealed and subcloned into a blunt pBluescript SK (+) plasmid. This plasmid was subsequently digested with NotI and KpnI and the T2A-NLS fragment was cloned into the tdTOM plasmid. tdTOM was amplified by PCR from the initial plasmid (primers 7 and 8) adding KpnI and FseI sites and subcloned into the donor vector after the 3xNLS sequence. An FNF cassette containing two FRT sites, and the NeoR/KanR gene under the control of the PGK promoter was amplified by PCR (primers 9 and 10) adding FseI and PacI sites. This fragment was subcloned into the donor vector.

The highest scoring sgRNA sequence to target the *Myog* STOP codon region was determined using the guide design tool from crispr.mit.edu (Zhang Lab). Primers containing this targeting sequence (primers 11 and 12) were annealed and subcloned into the pU6-(BbsI) CBh-Cas9-T2A-mCherry vector (Addgene #64324) digested with BbsI.

### Targeting of mouse embryonic stem cells

The Myog-ntdTOM donor construct (linearized by PvuI digestion) and the pU6 vector were electroporated in *C57BL/6J* mouse embryonic stem cells. Following G418 (300 μg/mL) selection, positive clones were determined by PCR using primers 13, 14 and 15, yielding a 1.7 kb band for the WT and 1.2 kb for the mutant. Two positive clones were expanded and 8-10 embryonic stem cells were injected into *BALB/c* blastocysts to generate chimeric mice in the Mouse Genetics Engineering Facility at the Institut Pasteur. Germline transmission was verified by PCR and F1 mice were crossed to *Tg(ACTFLPe)9205Dym* [38] animals to excise the FNF cassette. Excision of the FNF cassette and presence of the *Myog*^*ntdTom*^ allele was verified by PCR using primers 16, 17 and 18 (Flp-recombined *Myog*^*ntdTom*^ allele, 236 bp and WT allele 600 bp, **Figure S1**) and these primers were subsequently used for genotyping. F2 animals were backcrossed to *C57BL/6* animals and *Myog*^*ntdTom/+*^; *Tg(ACTFLPe)*^*+/+*^ animals were selected for further characterisation.

### Isolation and culture of MuSCs

Fetal and adult muscles were dissected and minced in ice-cold DMEM as described in [39]. Samples were then incubated in DMEM, 0.08% Collagenase D (Sigma, 11088882001), 0.2% Trypsin (ThermoFisher, 15090) and 10 µg/ml of DNAseI (Sigma, 11284932) for 25 min at 37°C under gentle agitation for 5 rounds of digestion. After each round, samples were allowed to sediment for 5 min, the supernatant was collected in 4ml of Fetal Bovine Serum (FBS) on ice and fresh digestion buffer was added to the remaining muscle pellet. The collected supernatants were centrifuged for 15 min at 550 g at 4°C, resuspended in DMEM 2% FBS and filtered through a 40µm strainer (Corning, 352235) before cell sorting. Cells were isolated based on size, granulosity and GFP or tdTOM fluorescence on using an Aria III (BD Biosciences) flow cytometer. Cells were collected directly in MuSC growth media (38.5% DMEM (Fisher Scientific, 31966047), 38.5% F12 (Fisher Scientific, 31765035), 20% FBS (ThermoFisher, 10270), 2% Ultroser (Pall, 15950-017), 1% Penicillin/Streptomycin (GIBCO, 15140-122)).

Matrigel^®^ (1 mg/ml, Corning, 354248) coated dishes (30 min at 37°C) were used to culture MuSCs in growth media at 3% O_2_, 5% CO_2_, 37°C for the indicated times.

For immunostaining, cells were fixed in 4% paraformaldehyde (PFA, Electron Microscopy Sciences, 15710) for 15 min at RT, permeabilised in 0.5% Triton X-100 (Merck, T8787) for 5 min at RT and blocked with 10% goat serum (GIBCO). Cells were incubated with the indicated primary antibodies in PBS 2% FBS buffer overnight following by 45 min incubation with secondary antibodies and 1 μg/ml Hoechst (ThermoFisher, H1399).

### Embryo immunofluorescence

For tissue immunofluorescence, embryos were collected in PBS and fixed in 4%PFA 0.1% Triton X-100 in PBS for 2h at 4°C. After 3 PBS washes, embryos were cryopreserved in 30% sucrose in PBS and embedded in OCT tissue freezing media (Leica, 14020108926) for cryosectioning. Cryosections were allowed to dry for 30 min at room temperature and washed once with PBS. Tissue samples were blocked in 3% BSA, 10% goat serum, 0.5% Triton X-100 for 1h at room temperature. Primary antibodies were diluted in blocking solution and incubated overnight at 4°C. After three washes PBST (PBS 0.1% Tween20 (Sigma Aldrich, P1379)), secondary antibodies were diluted in blocking solution and incubated for 45 min at room temperature. Finally, samples were incubated with 1 μg/ml Hoechst 33342 for 5 min at room temperature to visualize nuclei, washed three times in PBS and mounted in 70% glycerol in PBS for imaging.

For whole mount immunofluorescence embryos were collected in PBS and fixed in 4% PFA 0.1% Triton X-100 for 2h at 4°C. After two PBS washes, samples were dehydrated in 50% methanol in PBS and kept in 100% methanol at −20°C until used. Samples were rehydrated in PBS and incubated in blocking buffer (10% goat serum, 10% BSA, 0.5% TritonX-100 in 1X PBS) for 1 hr at RT in 2 ml Eppendorff tubes. Embryos were then incubated with primary antibodies in the blocking buffer for 5-7 days at 4°C with rocking. Embryos were washed extensively for 2-4 h in PBST and incubated in Fab’ secondary antibodies for 2 days at 4°C with rocking. Embryos were washed as above, dehydrated in 50% Methanol in PBS, twice in 100% Methanol and then cleared with BABB and mounted for imaging [40].

### Adult muscle injury, histology and immunofluorescence

Muscle injury was done as described previously [22]. Mice were anesthetized with 0.5% Imalgene/2% Rompun and the TA muscle was injected with 50 mL of Cardiotoxin (10mM; Latoxan, L8102) diluted in 0.9% NaCl.

Injured TA muscles were fixed upon harvesting in 4% PFA for 2h at 4°C, washed with PBS and equilibrated with 30% sucrose in PBS overnight. Samples were mounted in OCT tissue freezing media and cryosectioned between 8-12 μm. When endogenous tdTOM was scored, cryosections were rehydrated in PBS and counterstained with Hoechst 33342.

In case of MYOG plus tdTOM detection, tissue sections were processed for histology as described [41]. Briefly, sections were post-fixed in 4% PFA for 10 min at RT and washed with PBS prior to immunostaining. Heat-induced epitope retrieval was performed in a citrate solution pH 6.0 during 6 min in a pressure cooker. Sections were then incubated with 30% H2O_2_ for 5 min at RT. Samples were then permeabilised with 0.2% Triton-X100, washed in PBS and blocked in blocking buffer (10% goat serum, 10% BSA, 0.5% TritonX-100 in 1X PBS). Primary antibodies against MYOG and DsRED (recognizing tdTOM) were incubated overnight at 4°C. After washing with PBST, sections were incubated with appropriate secondary antibodies and 1 μg/ml Hoechst 33342 in blocking buffer for 45 min at RT.

**Table 1.**
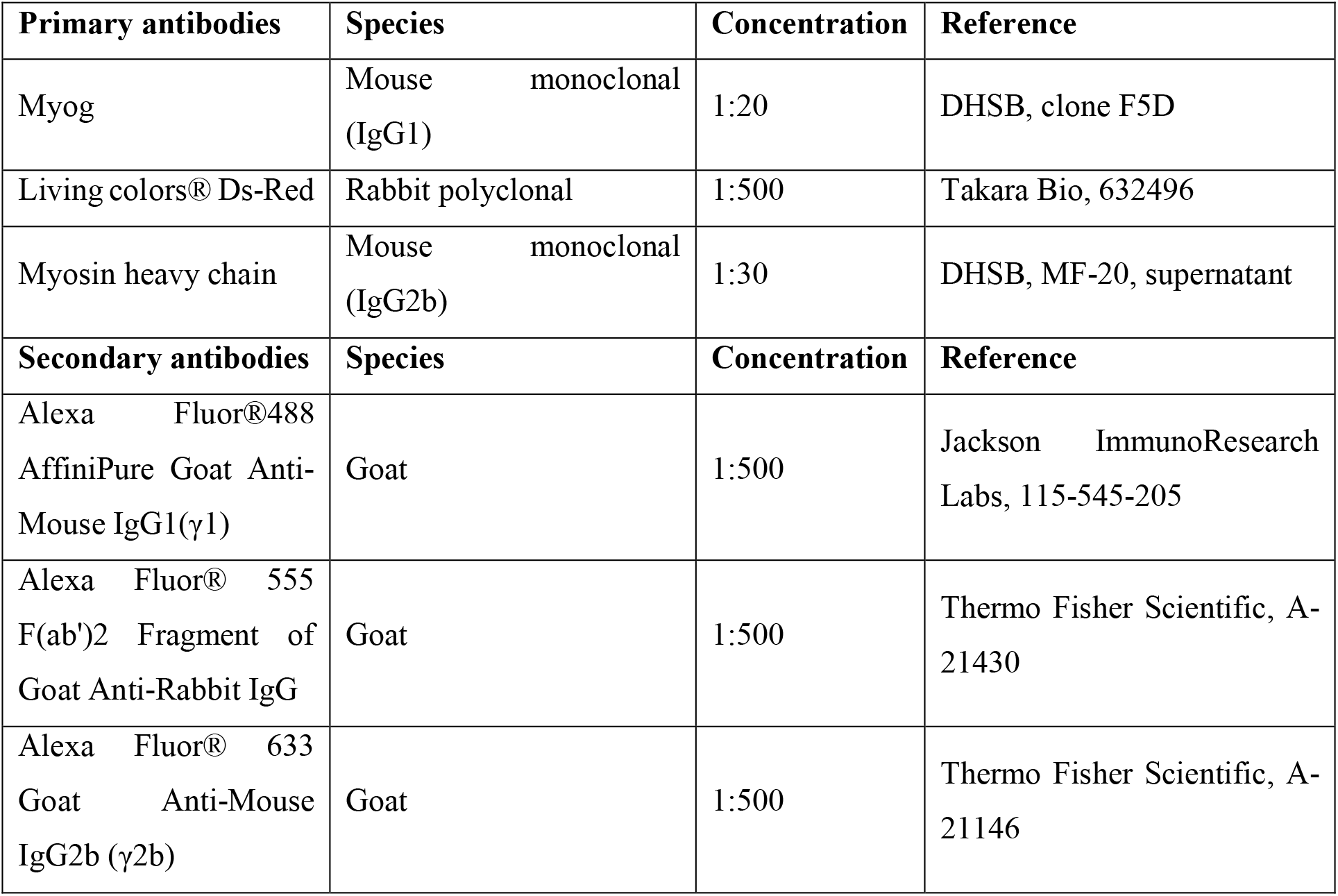
Antibodies used for immunostaining.

### Western Blot

Embryos were collected in ice cold PBS and subsequently snap frozen in dry ice. Embryonic tissues were ground to a fine powder using a mortar and pestle on dry ice and lysed in RIPA buffer (150M NaCl, 50mM Tris pH8, 5mM EDTA, 1% NP-40 (Sigma, I8896), 0.5% sodium deoxycholate, 0.1% SDS supplemented with 1X proteases (Sigma, S8820) and phosphatases (Roche, 4906845001) inhibitors). 15 ug of protein extracts were run on a 4%–12% Bis-Tris Gel NuPAGE (Invitrogen, NP0322) and transferred on a PVDF Amersham Hybond-P transfer membrane (GE Healhcare, RPN303F). The membrane was blocked with 5% milk in Tris-Buffer Saline 0.2% Tween (Sigma, P9416) (TBS-T) for 1h at room temperature and probed with specific primary antibodies overnight at 4°C. After three washes in TBS-T, the membrane was incubated with HRP or fluorophore-conjugated secondary antibodies and revealed by chemiluminescence (Pierce ECL2 western blotting substrate, Thermo Scientific, 80196) or fluorescence (Bio-Rad, Chemidoc MP).

**Table 2.** Antibodies used for Western Blot.

| Primary antibodies | Species | Concentration | Reference |
| --- | --- | --- | --- |
| Myog | Mouse monoclonal (IgG1) | 1:500 | DHSB, clone F5D |
| Living colors® Ds-Red | Rabbit polyclonal | 1:1000 | Takara Bio, 632496 |
| Secondary antibodies | Species | Concentration | Reference |
| Mouse-HRP | Goat polyclonal | 1:5000 | Pierce, 31430 |
| Rabbit-HRP | Goat polyclonal | 1:5000 | Pierce, 31460 |
| GAPDH-Rhodamine | Human IgG Fab fragment | 1:5000 | Bio-Rad, 12004168 |

### RNA extraction

RNA from cells isolated by FACS was extracted using a Trizol-based kit (Zymo Research, R2061) and reverse transcribed using SuperScriptIII (Invitrogen, 18080093). RT-qPCR to assess for mRNA relative expression was performed with SYBR green master mix (Roche, 04913914001) in Applied biosciences machine. Data analysis was performed using the 2^-ΔΔCT^ method [42] and mRNA expression was normalized with *Rpl13*.

### *In vitro* videomicroscopy

MuSC were plated on a microscopy culture chamber (IBIDI, 80826) and cultured in growth media supplemented as above. The plate was incubated at 37°C, 5% CO_2_, and 3% O_2_ in a Pecon incubation chamber. A Zeiss Observer.Z1 connected to a Plan-Apochromat 20x/0.8 M27 objective and Hamamatsu Orca Flash 4 camera piloted with Zen software (Carl Zeiss) was used.

### Static imaging

The following systems were used for image acquisition: Zeiss SteREO Discovery V20 for macroscopic observations of whole embryos and Zeiss LSM800 or LSM700 laser-scanning confocal microscopes for tissue sections and whole mount immunostaining of cleared embryos. End point in vitro culture samples were imaged with a Zeiss Observer.Z1.

### Intravital microscopy

Intravital imaging of *Pax7*^*CreERT2*^; *R26*^*YFP*^; *Myog*^*ntdTom*^ mice at different timepoints during regeneration was performed on an upright Nikon NiE A1R MP microscope piloted with NIS software (Nikon). The microscope was equipped with a 25x NA 1.1 PlanApo LambdaS objective, GaAsP PMT detectors and a Spectra-Physics Insight Deepsee laser. Laser frequency was tuned to 960 nm to allow simultaneous excitation of YFP and tdTOM fluorophores.

For image acquisition, the skin over the upper hindlimb was shaved and incised to expose approximately 1 cm^2^ of the muscle and imaged directly. During the imaging period mice were anesthetised with 1.5% isofluorane and maintained in an incubation chamber at 37°C.

### Image analysis

Cell tracking was performed using the Manual Tracking feature of the TrackMate plug-in [43] in Fiji [44]. ZEN software (Carl Zeiss), Fiji [44] and Imaris (Bitplane) were used for image analysis. Figures were assembled in Adobe Photoshop and Illustrator (Adobe Systems).

### Data analysis and statistics

Data analysis and statistics were performed using R [45] and figures were produced using the package ggplot2 [46]. For comparison between two groups, two tailed paired and unpaired Student’s t-tests were performed to calculate P values and to determine statistically significant differences (see Figure Legends).

**Table 3.**
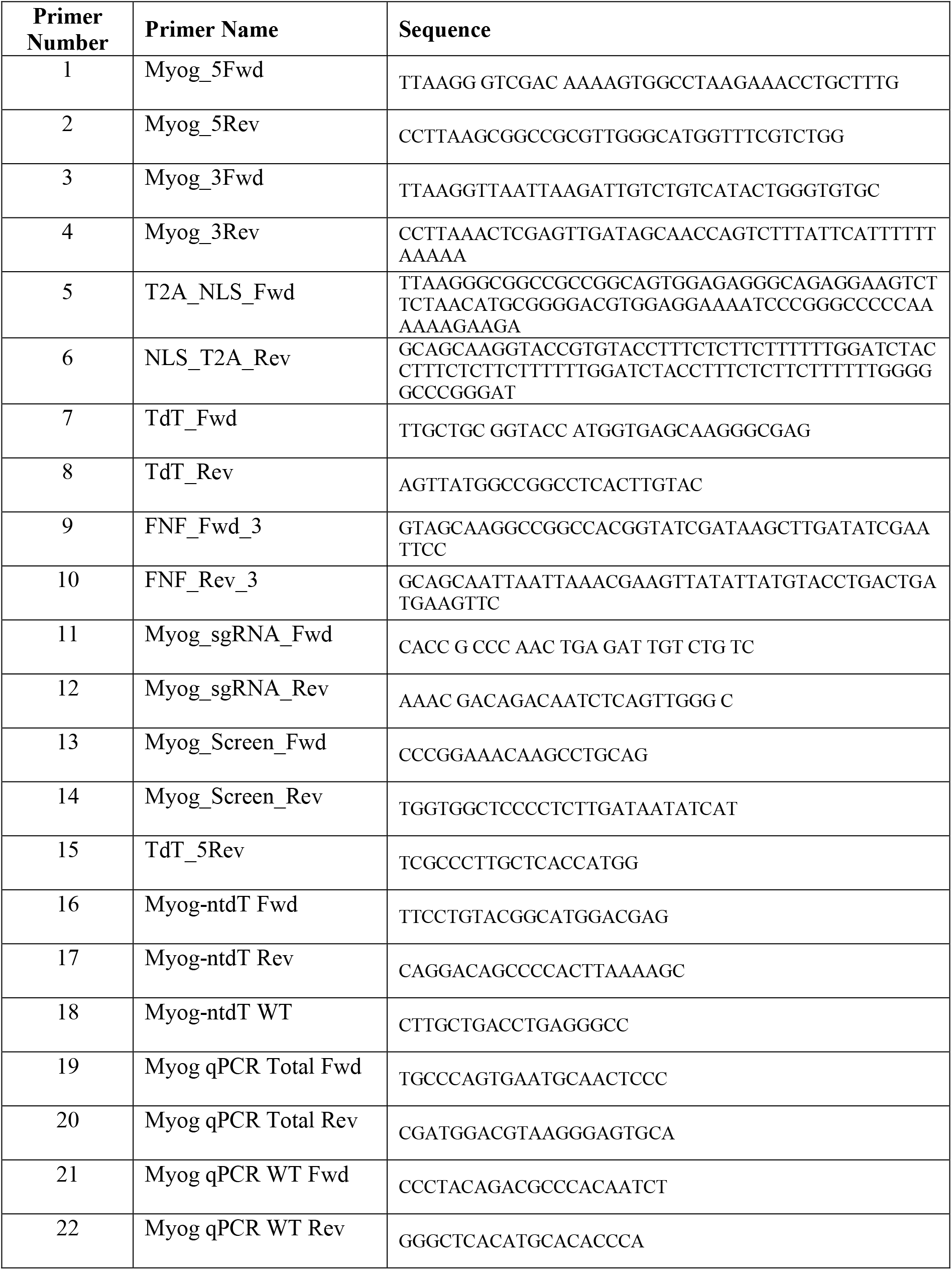
Primer sequences.

## RESULTS

### Generation and characterisation of a *Myog*^*ntdTom*^ mouse

Using the CRISPR-Cas9 system, a sgRNA was designed to target the region of the STOP codon of the *Myog* gene for homologous recombination. The recombination template consisted of two homology arms corresponding to *Myog* sequences flanking the STOP codon, a *tdTom* coding sequence and a Neo resistance cassette flanked by *frt* sites (**Figure 1A**). The tdTOM protein was preceded by a T2A peptide sequence [36] to allow cleavage from the MYOG protein following translation, and a triple NLS sequence [37] to ensure nuclear localisation.

**Figure 1.**
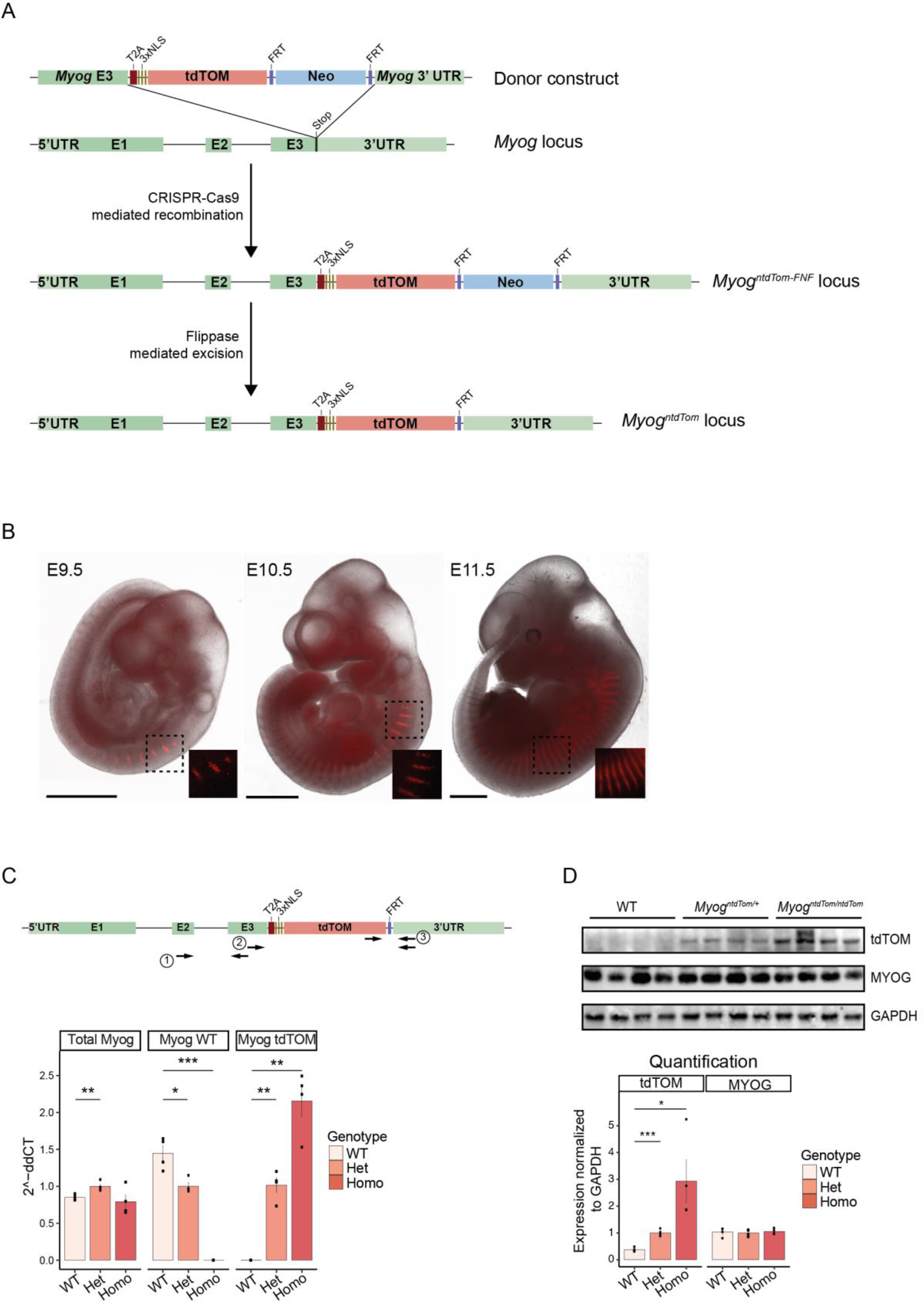
Generation of a *Myog* knock-in mouse line. A. Scheme depicting the endogenous *Myog* locus, the donor construct, and the result of the CRISPR-Cas9-mediated recombination in mouse embryonic stem cells. First generation *Myog*^*ntdTom-FNF*^ mice were then crossed with a *Tg(ACTFLPe)*^*+/+*^ deleter strain to excise the FNF cassette. B. Endogenous fluorescence from *Myog*^*ntdTom/+*^ embryos at different stages. An overlay between the brightfield and fluorescent images is shown. Scale bar, 1000 μm. C. Scheme showing the primer pairs amplifying the wild-type allele (2), the *ntdTom* allele (3) and both (1) in the targeted *Myog* locus. RT-qPCR analysis of the levels of total *Myog* mRNA, the wild-type allele and the *tdTom* allele specifically from E14.5 *Myog*^*+/+*^, *Myog*^*tdT/+*^ *Myog*^*tdT/tdT*^ embryos. n=4 embryos per genotype. Data represents mean ± s.d. Two-tailed unpaired Student’s t-test; *** p-value < 0.005, ** p-value = 0.0005 to 0.01, * p-value = 0.01 to 0.05. D. Western blot assessing the levels of MYOG and tdTOM proteins from E14.5 *Myog*^*+/+*^, *Myog*^*ntdTom/+*^ and *Myog*^*ntdTom/ntdTom*^ embryos (n=4 embryos per genotype). Bar graph shows the quantification of protein expression levels normalized to GAPDH. Data represents mean ± s.d. Two-tailed unpaired Student’s t-test; *** p-value < 0.005, * p-value = 0.01 to 0.05.

First, we evaluated the endogenous tdTOM fluorescence in heterozygous *Myog*^*ntdTom/+*^ embryos between E9.75 and E11.5 at the level of the somites, i.e. transient embryonic structures arising from the segmentation of the paraxial mesoderm. Endogenous tdTOM fluorescence followed a similar pattern to that described for *Myog* transcripts (Cheng et al., 1992) (**Figure 1B**), with expression levels being lower in the caudal (more recently formed) somites as expected.

We then collected tissue samples from E14.5 fetuses and performed RT-qPCR and Western blot analysis to confirm that *Myog* mRNA and protein levels were similar in wild-type, heterozygous and homozygous animals. Primer pairs were designed to amplify specifically the wild-type allele or the *tdTom* allele, and one primer set amplified both (**Figure 1C**). This analysis showed that *Myog* heterozygous and homozygous knock-in (KI) embryos expressed similar levels of total *Myog* mRNA, and confirmed that no *Myog* wild-type transcript could be detected in the homozygous embryos. As expected, the levels of *Myog*^*ntdTom*^ mRNA were the highest in homozygous samples, decreased to roughly 50% in the case of the heterozygous, and was not detected in wild-type embryos (**Figure 1C**). At the protein level, we noted similar expression levels of MYOG in embryos from all three genotypes, whereas the tdTOM protein was absent in wild-type samples (**Figure 1D**). Therefore, we conclude that MYOG protein was generated from transcripts that originated from both alleles.

To investigate the expression of the targeted allele with higher resolution, we assessed the temporal expression dynamics of MYOG and tdTOM proteins by whole mount immunostaining at E10.5 and compared the expression of MYOG and tdTOM in wild-type, heterozygous and homozygous embryos (**Figure 2A, Additional files 1-3**). We confirmed that tdTOM followed the expression pattern of MYOG in the epaxial and hypaxial domains of all somites, indicating that both proteins have similar spatiotemporal expression dynamics. To assess the co-expression of MYOG and tdTOM at the single cell level in heterozygous embryos, cryosections at the level of extraocular, tongue and limb muscles were examined during primary (E12.5) and secondary (E14.5) myogenesis when small oligo-nucleated and larger multi-nucleated myofibres are generated, respectively. Quantification of protein expression in different muscles confirmed an average co-localisation of both proteins in 97% and 95% of the cells at E12.5 (**Figure 2B**) and E14.5 (**Figure S2A**), respectively.

**Figure 2.**
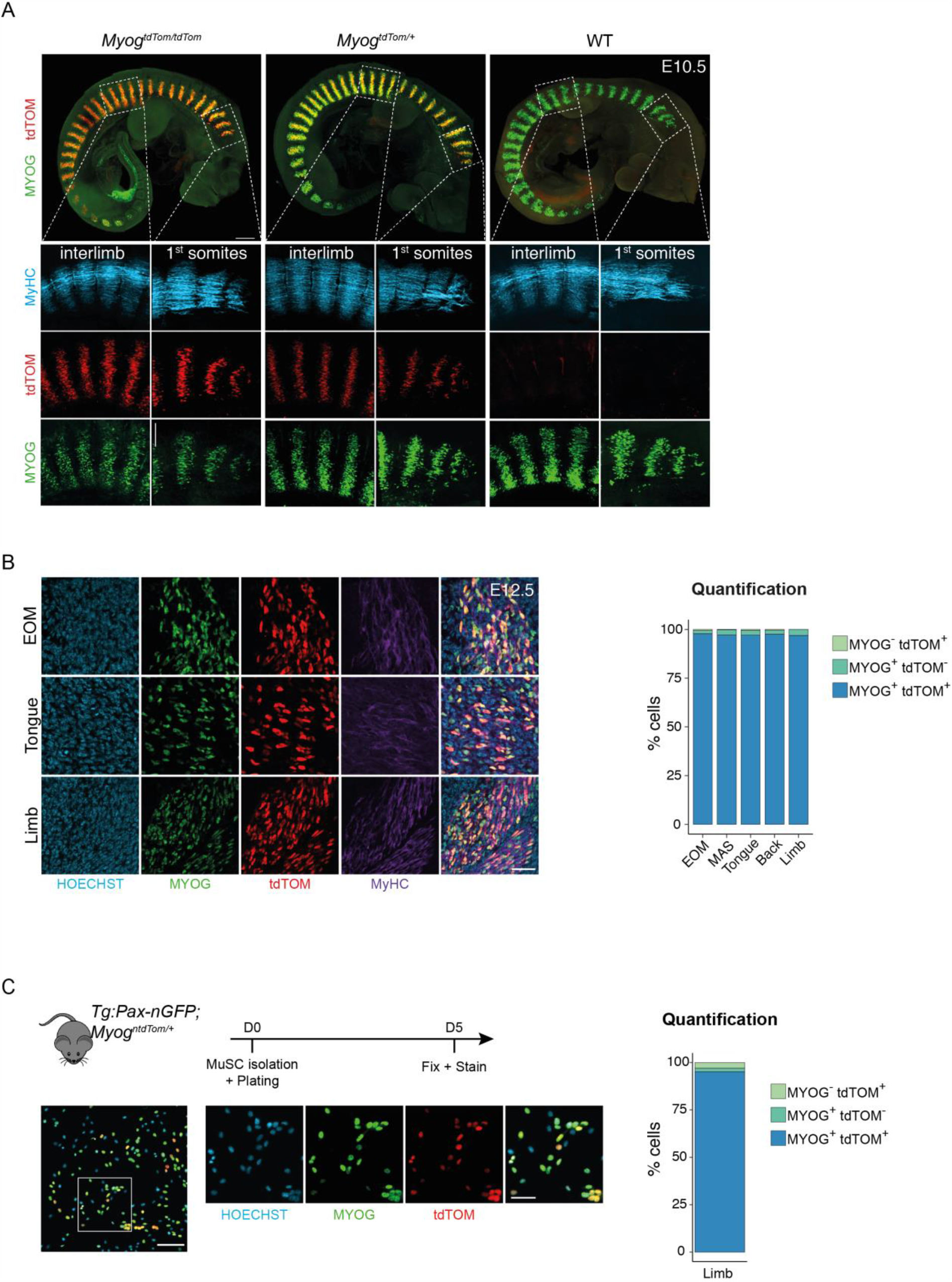
tdTOM expression recapitulates endogenous *Myog* expression. A. Whole mount immunofluorescence from *Myog*^*+/+*^, *Myog*^*ntdTom/+*^ and *Myog*^*ntdTom/ntdTom*^ embryos at E10.5 stained for tdTOM, MYOG and MyHC proteins. Scale bar, 300 μm on upper panels and 100 μm on lower insets. B. Immunofluorescence of extraocular, tongue and limb muscle cryosections of *Myog*^*ntdTom/+*^ embryos at E12.5. The bar graph shows the quantification of tdTOM and MYOG positive cells. n=3 embryos, >100 cells per muscle per embryo were counted. Scale bar, 40 μm. C. MuSCs were isolated from limb muscles of *Tg:Pax7-nGFP; Myog*^*ntdTom*^ animals and plated for *in vitro* differentiation for 5 days. Bar graph shows the quantification of tdTOM and MYOG positive cells (n=3, >200 cells per animal counted). Scale bar, 100 μm on left image, 50 μm on right inset.

To assess the fidelity of the reporter mouse in adult myoblasts, *Myog*^*ntdTom*^ animals were crossed with *Tg:Pax7-nGFP* mice, where GFP marks all MuSCs [15]. MuSCs were isolated by FACS based on GFP expression, then differentiated *in vitro* for 5 days. In agreement with our results in the embryo, MYOG and tdTOM expression co-localised in about 95% of the cells (**Figure 2C**). Additionally, no significant differences were observed in total *Myog* RNA levels between wild-type, heterozygous and homozygous animals (**Figure S2B**).

As indicated above, the KI strategy was designed to be non-disruptive and allow normal MYOG protein expression from the recombined alleles. Given that *Myog* null mice are lethal at birth, and our *Myog*^*ntdTom/ntdTom*^ knock-in mice are viable, we propose that sufficient levels of MYOG are produced from the targeted allele. Nonetheless, a decrease in MYOG intensity was detected by immunofluorescence in homozygous *Myog*^*ntdTom/ntdTom*^ embryos and *in vitro* myoblast cultures from homozygous animals (**Figure 2A, Figure S2C**). As this decrease was not observed in heterozygous samples, we decided to use heterozygous animals in our subsequent experiments.

In summary, tdTOM faithfully recapitulates the expression of MYOG protein in embryonic and adult muscle, and its insertion at the *Myog* locus does not impair significantly the expression of this gene at the mRNA and protein level.

### *Myog*^*ntdTom*^ mice allow isolation of differentiating myoblasts at different stages

To assess the expression of tdTOM in homeostatic conditions by flow cytometry, we isolated the mononuclear population from limb muscles of *Myog*^*ntdTom/+*^ mice at fetal (embryonic day (E) 18.5), postnatal (postnatal day (p) 21) and adult (10 weeks) stages. tdTOM fluorescent cells were detected at fetal and early postnatal stages where myogenesis was still taking place. In adult muscles in homeostasis, the majority of MuSCs are quiescent and therefore no MYOG^+^ mono-nucleated cells are detectable (Gattazzo et al., 2020). As expected, virtually no tdTOM^+^ cells were detected in muscles of adult *Myog*^*ntdTom/+*^ animals (**Figure 3A**).

**Figure 3.**
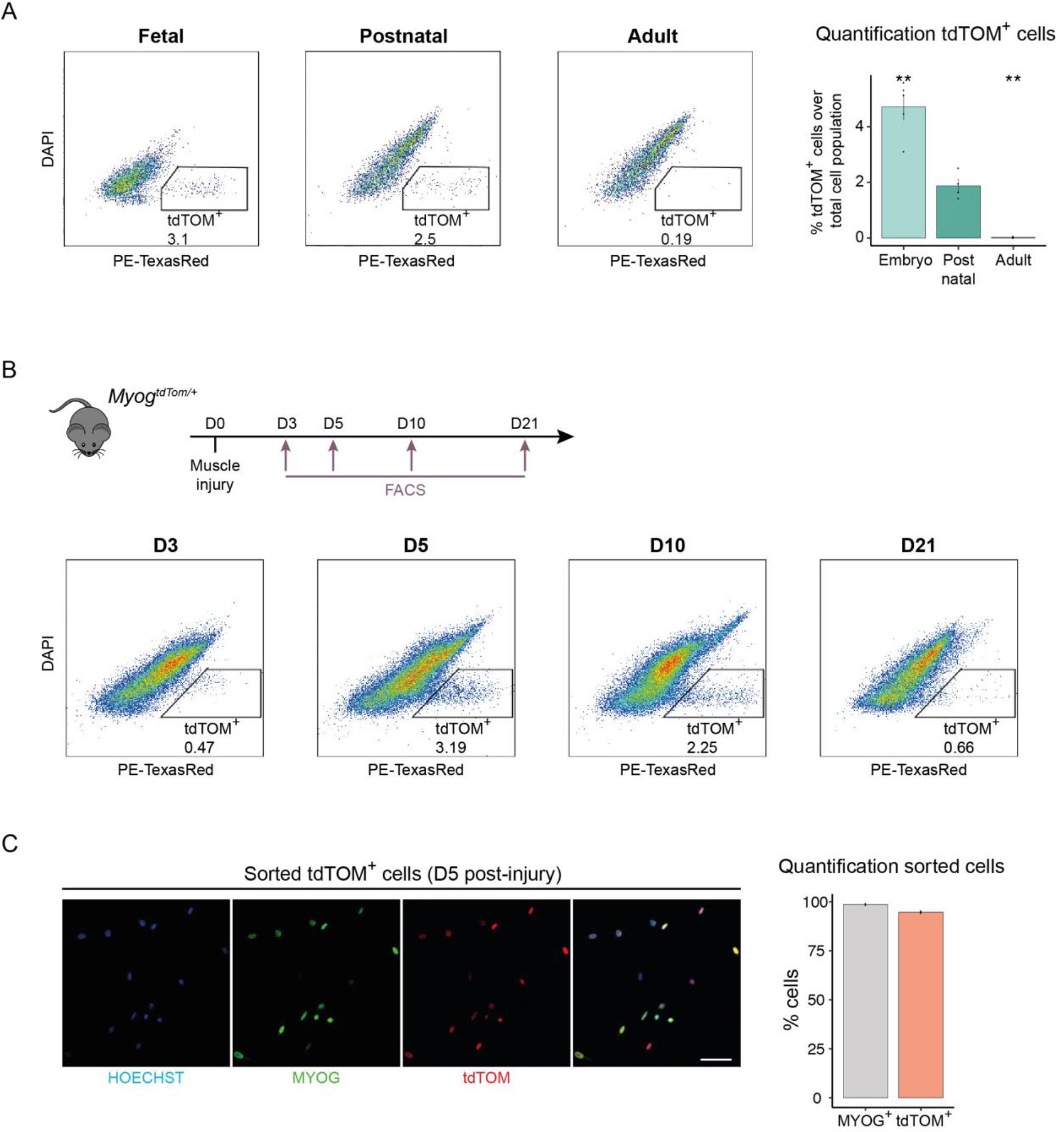
*Myog*^+^ cells can be isolated based on tdTOM fluorescence. A. FACS profiles of the mononuclear cell fraction isolated from limb muscles of E18.5, p21 and p70 of *Myog*^*ntdTom/+*^ animals. Bar graph shows the percentage of tdTOM^+^ cells at the different stages. n>4 animals per condition. Data represents mean ± s.d. Two-tailed unpaired Student’s t-test; ** p-value = 0.0005 to 0.01. B. FACS profiles of the mononuclear cell fraction isolated from cardiotoxin injured TA muscles from *Tg:Pax7-nGFP; Myog*^*ntdTom/+*^ mice at different timepoints (D3, D5, D10, D21 post-injury). C. Immunostaining of the tdTOM^+^ cell fraction at 5 DPI isolated as in (B) for MYOG (3h post plating). Bar graph shows the quantification of tdTOM and MYOG positive cells (n=3, >200 cells per animal counted). Data represent mean ± s.d. Scale bar, 50 μm.

To determine if tdTOM followed the expression dynamics of MYOG during adult muscle regeneration, we performed an injury of the *Tibialis anterior* (TA) muscle of *Tg:Pax7-nGFP; Myog*^*ntdTom/+*^ mice by intramuscular injection of the snake venom toxin cardiotoxin [47]. We next performed FACS analysis to determine whether the tdTOM^+^ mononucleated fraction could be isolated following tissue injury. As expected, only a few tdTOM^+^ cells were detected at 3 days post-injury, when myogenic cells are known to be maximally proliferating [23,24,48] (**Figure 3B**). tdTOM^+^ cells were most abundant at 5 and 10 days post injury, corresponding to the differentiation shift of the transiently amplifying myoblast population. As the major features of the regeneration process are completed by 3-4 weeks, the proportion of tdTOM^+^ cells decreased by 21 days post-injury, corresponding to the progressive return to quiescence of the myogenic population (**Figure 3B**). Finally, to verify whether the tdTOM^+^ cells isolated by FACS corresponded to myoblasts that expressed MYOG, we isolated the tdTOM^+^ population from regenerating TA muscle at 5 days post-injury. Fixation of cells immediately after sorting and staining for MYOG and tdTOM showed that 95% of isolated cells were positive for MYOG (**Figure 3C**), thereby confirming that tdTOM followed the expression dynamics of MYOG during adult muscle regeneration and that its expression allows isolation of MYOG^+^ cells by FACS after injury.

Taken together, our results show that the *Myog*^*ntdTom*^ KI mouse allows efficient isolation of the MYOG^+^ population at different stages during development as well as from regenerating muscle.

### Dynamics of *Myog* expression during terminal differentiation

To assess if *Myog*-expressing myoblasts can execute cell division, we took advantage of the tdTOM reporter to monitor *Myog* expression by live videomicroscopy of primary myoblasts *in vitro*. MuSCs from adult *Tg:Pax7-nGFP; Myog*^*ntdTom/+*^ mice were isolated by FACS based on GFP fluorescence and plated for *in vitro* differentiation. After 3 days of *in vitro* culture, live-imaging was started and images were acquired every 12 min for 48 hours (**Figure 4A, Additional File 4)**.

**Figure 4.**
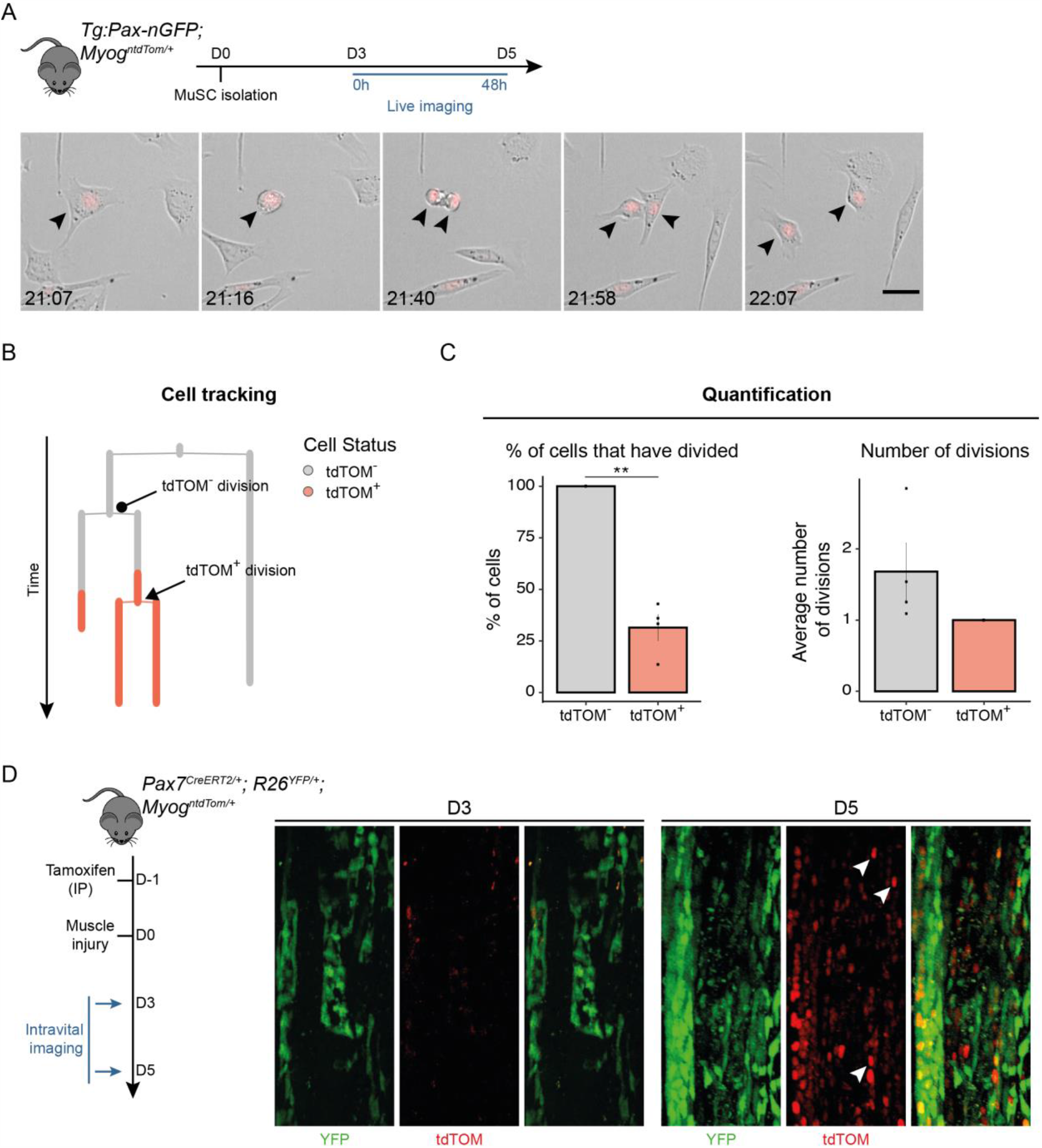
*Myog*^+^ cells can undergo cell division. A. MuSCs were isolated based on GFP fluorescence from *Tg:Pax7-nGFP*^*Tg/+*^; *Myog*^*ntdTom/+*^ mice. Cells were plated for 3 days before initiating live imaging. Black arrows show a tdTOM^+^ cell dividing and its daughter cells. Images were acquired every 9 min. Scale bar, 25 μm. B. Representative tracking output from experiment in (A). C. Left bar graph shows the percentage of tdTOM^-^ and tdTOM^+^ cells that have undergone at least one cell division. Right bar graph indicates the average number of divisions that tdTOM^-^ and tdTOM^+^ underwent during the tracking period (n=100 cells tracked). Data represent mean +/- s.d. Two-tailed unpaired Student’s t-test; ** p-value = 0.005 to 0.01. D. Recombination of *Pax7*-expressing cells in *Pax7*^*CreERT2*^; *R26*^*YFP*^; *Myog*^*ntdTom*^ reporter mice was induced 1 day before injury of the upper hindlimb muscle. Images were acquired by intravital imaging at 2 timepoints during muscle regeneration (1 mouse/timepoint). White arrows indicate double positive YFP and tdTOM cells. Scale bar, 100 μm.

By manually tracking individual cells and monitoring their differentiation status based on tdTOM fluorescence, we observed that up to 35% of MYOG^+^ cells underwent cell division during the imaging period (**Figure 4B-C**). Having established that MYOG^+^ cells remain competent for cell-division, we sought to quantify the number of divisions that they performed. Amongst all the MYOG^+^ cells tracked, none divided more than once. In contrast, all MYOG^-^ cells divided during the imaging period and performed 1.5 divisions on average during this time (**Figure 4C**). Therefore, using the *Myog*^*ntdTom*^ reporter mouse, we show that a significant proportion of MYOG^+^ cells can undergo one more cell division when tracking tdTOM expression.

Finally, we assessed the potential of the *Myog*^*ntdTom*^ mouse to monitor tissue regeneration by intravital imaging. *Pax7*^*CreERT2/+*^; *R26*^*YFP/+*^; *Myog*^*ntdTom/+*^ mice were used to permanently label *Pax7*-expressing cells and their progeny upon tamoxifen administration and simultaneously trace the differentiated fraction by following tdTOM expression. We induced muscle injury by cardiotoxin injection in the upper hind limb and monitored the regeneration process by live intravital imaging at different time points (**Figure 4D**). Few MYOG^+^ cells were detected at 3 days post-injury, whereas extensive proliferation of YFP^+^ myogenic progenitor cells was taking place (**Figure 4D**, left panels). Two days later, the population of tdTOM^+^ myoblasts had significantly expanded and tdTOM^+^ cells could be observed throughout the regenerating area (**Figure 4D**, right panels, white arrows), recapitulating the results of our flow cytometry analysis.

Taken together, these experiments demonstrate that tdTOM expression is robust enough to monitor Myog^+^ cells by *in vitro* and intravital imaging.

## DISCUSSION

*Myog* is a critical regulator of myoblast differentiation and fusion, being an essential factor for embryonic muscle development. In the present study we generated and characterised a novel mouse line to fluorescently label MYOG^+^ cells by co-expression of the robust nuclear localised tdTOM protein from the endogenous *Myog* locus.

To characterise the properties of tdTOM^+^ cells, we assessed its co-localization with MYOG during embryonic and fetal development, in adult primary myogenic cells *in vitro*, and during adult muscle regeneration *in vivo*. Given that in all conditions virtually all cells were positive for both markers (>95% cells), we conclude that tdTOM reliably labels MYOG^+^ cells from development to adulthood. Furthermore, total *Myog* levels from *Myog*^*ntdTom/+*^ animals were comparable to that of the WT, and homozygous *Myog*^*ntdTom/ntdTom*^ mice are viable. Moreover, expression of tdTOM allowed us to isolate the MYOG^+^ population from developing embryos as well as adult regenerating muscle indicating the utility of this reporter mouse in isolating living differentiating myoblasts that were previously inaccessible for direct investigation.

In addition, studies on the cell cycle dynamics of *Myog*^+^ cells have been hampered by the lack of a fluorescent reporter. Here, by means of live microscopy and single-cell tracking of differentiating primary myoblasts, we demonstrated that about one third of MYOG^+^ cells can divide *in vitro* and undergo a maximum of one additional cell division during the tracking period. Therefore, the majority of cells that express detectable levels of MYOG exit the cell cycle.

Several studies have performed intravital imaging of muscle tissue [24,49–51], however, only two of them dynamically monitored the process of muscle regeneration [24,49]. These two studies focused on the progenitor population by labelling PAX7^+^ cells, but they did not report on the dynamics of differentiation. Here, we carried out proof of concept experiments by intravital imaging of adult regenerating muscle and showed that tdTOM fluorescence is sufficient to follow MYOG^+^ cells throughout the regeneration process.

## CONCLUSION

In this study, we describe the creation of a new mouse line where tdTOM is expressed from the endogenous *Myog* locus. tdTOM recapitulates Myog expression during embryonic development and adult muscle regeneration and can be used to isolate this population by flow cytometry. Additionally, heterozygous tdTOM expression is sufficient for monitoring *Myog* dynamics by *in vivo* intravital imaging. Therefore, the *Myog*^*ntdTom*^ line can be of great benefit to study the dynamics of lineage progression of muscle progenitors in embryonic and adult stages.

## ADDITIONAL FILES

## Abbreviations

Myog: Myogenin
MFR: Myogenic Regulatory Factor
MuSC: Muscle Stem Cell
BrdU: 5-bromo-2’-deoxyuridine
sgRNA: Single Guide RNA
CRISPR: Clustered Regularly Interspaced Short Palindromic Repeats
NLS: Nuclear Localisation Sequence
GFP: Green Fluorescent Protein
RT-qPCR: Real-Time Quantitative Polymerase Chain Reaction
KI: Knock-in
FACS: Fluorescence Assisted Cell Sorting
TA: Tibialis Anterior
YFP: Yellow Fluorescent Protein

## Acknowledgements

We gratefully acknowledge S. Paisant for help with the maintenance of mouse embryonic stem cell lines, the Institut Pasteur Mouse Genetics Engineering Platform, the UtechS Photonic BioImaging (Imagopole), at Institut Pasteur, supported by the French National Research Agency (France BioImaging; ANR-10–INSB–04; Investments for the Future), the Center for Translational Science (CRT)-Cytometry and Biomarkers Unit of Technology and Service (CB UTechS) at Institut Pasteur and the Nikon Imaging Center at Institut Curie-CNRS for support in conducting this study.

## Availability of data and materials

All data generated and analysed during the study are available from the corresponding author on a reasonable request.

## Authors’ contributions

MBD designed, performed the experiments, and analysed the data. GC, DDG and FL performed some experiments. ST designed and supervised the project. MBD and ST interpreted data and wrote the manuscript. All authors edited the manuscript. All authors read and approved the final manuscript.

## Ethics approval

Animals were handled according to national and European Community guidelines and an ethics committee of the Institut Pasteur (CETEA) in France approved protocols (Protocol# 2015-0008).

## Consent for publication

All authors have read the final version of the manuscript and consented to its submission to Skeletal Muscle.

## FIGURES

**Figure S1.**
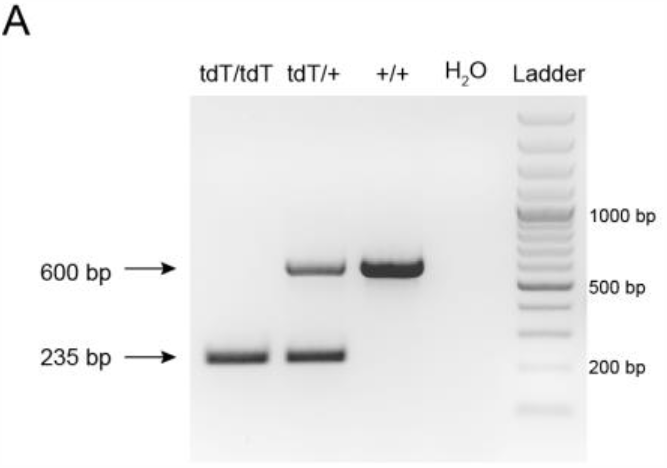
Genotyping of the *ntdTom* allele. **A**. Genotyping of ear-clip samples from *Myog*^*ntdTom/ntdTom*^, *Myog*^*ntdTom/+*^ and *Myog*^*+/+*^ animals. *Myog-ntdTom* allele was verified by PCR using primers 16, 17 and 18 (Flp-recombined *Myog-ntdTom* allele, 236 bp and WT allele 600 bp).

**Figure S2.**
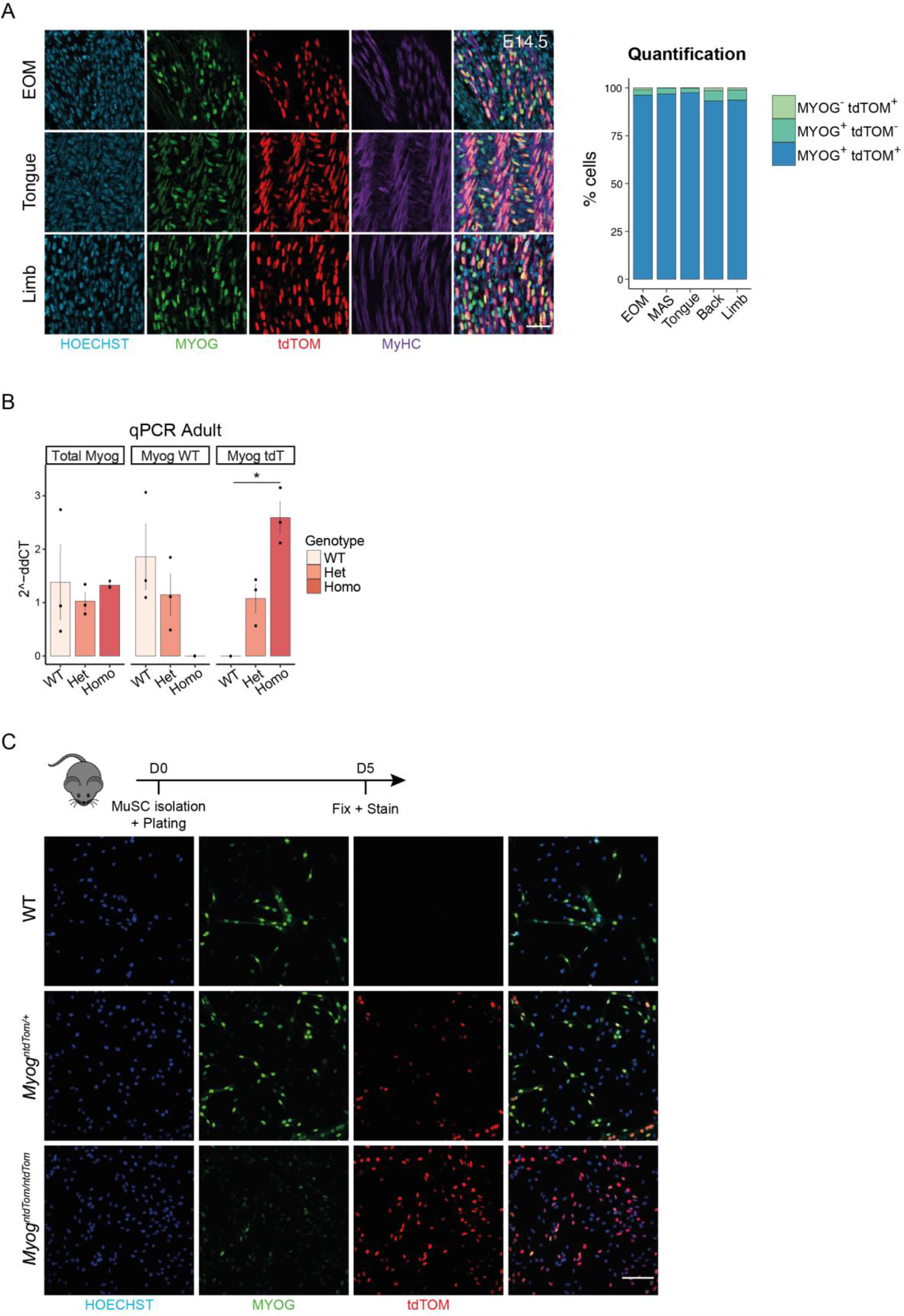
tdTOM expression recapitulates endogenous *Myog* expression. A. Immunofluorescence of extraocular, tongue and limb muscles of *Myog*^*ntdTom/+*^ embryo at E14.5. Bar graph shows the quantification of tdTOM and MYOG positive cells. n=3 embryos, >100 cells per muscle per embryo were counted. Scale bar, 40 μm. B. RT-qPCR assessing the levels of total *Myog* mRNA, the wild-type allele and the *tdTom* allele specifically from *Myog*^*+/+*^,*Myog*^*ntdTom/+*^ and *Myog*^*ntdTom/ntdTom*^ adult myoblasts using the primer set described in **Figure 1C**. n=3 animals per genotype. Data represents mean ± s.d. Two-tailed unpaired Student’s t-test; * p-value = 0.01 to 0.05. C. MuSCs from limb muscles were isolated from *Tg:Pax7-nGFP; Myog*^*+/+*^, *Tg:Pax7-nGFP; Myog*^*ntdTom/+*^ and *Tg:Pax7-nGFP; Myog*^*ntdTom/ntdTom*^ animals and plated for *in vitro* differentiation for 5 days. Cells were stained for MYOG and tdTOM proteins. Scale bar, 100 μm.

**Additional Files 1-3**. Related to Figure 2A. Whole mount immunofluorescence from *Myog*^*+/+*^, *Myog*^*ntdTom/+*^ and *Myog*^*ntdTom/ntdTom*^ embryos at E10.5 stained for tdTOM, MYOG and MyHC proteins.

**Additional File 4**. Related to figure 4A. *In vitro* division of a tdTOM^+^ cell.

